# Protein size and geometry govern mutational robustness

**DOI:** 10.64898/2026.08.20.745934

**Authors:** Srijita Acharya, Sucharita Dey

## Abstract

A fundamental question in structural biology centres around understanding protein evolution. Key to this process is mutational robustness, defined as the protein fold’s ability to absorb sequence changes without collapsing its structure. Here, we show that robustness is systematically shaped by simple features such as protein size, geometry, and oligomeric state. We used Foldseek-identified (structural) homologs to quantify family size across monomers and higher homo-oligomers. We found that proteins in larger families are consistently larger in size, more compact in atomic density, and less exposed to solvent. Strikingly, homo-oligomers occupy systematically larger families than monomers, revealing quaternary structure itself as a driver of mutational tolerance, not merely a functional supplement. This signature of robustness can be further linked to increasing functional complexity in proteins; those with adaptive, multifaceted biological roles belong to larger structural families than those with specific roles, thereby linking structural flexibility directly to evolutionary versatility. In short, simple yet overlooked features of protein geometry can explain mutational robustness and evolvability, offering a structural rationale for why certain protein families have diversified extensively while others remain in evolutionary stasis.

**eTOC blurb (short summary):** Why do some protein folds diversify into thousands of variants while others remain rare and rigid? This work shows that the answer lies in geometry: proteins with denser cores, larger size, and higher-order oligomeric assembly tolerate mutations more readily, occupy larger structural families, and support more versatile biological roles. This reveals that protein size, shape, and self-assembly, not just sequence, are fundamental drivers of evolvability.

## Introduction

Random genetic mutations are the primary driving force behind protein structure evolution, which are selected for their functional value. Some of these mutations, translated into amino acid substitutions in the protein structure, are most likely to cause loss of function or be deleterious, as they adversely affect structure, stability, and function. In contrast, some sets of mutations are retained and tolerated without significant disruption to protein structure and function (neutral mutations). In fact, the consequences of deleterious mutations are largely due to the disruption of a properly folded protein structure rather than to alterations in protein function.^1–5^ Proteins that can accommodate such mutational events are termed ‘robust’. Robustness refers to proteins’ ability to accumulate mutations with little to no change in structure, stability, or function. It is closely related to, but distinct from, evolvability, which describes changes in sequence that allow significant changes in structure and/or the introduction of a new function.^6,7^ While robustness allows for a neutral exploration of sequence space, evolvability allows the population to explore novel phenotypes, mediated by the accumulation of cryptic genetic variation.^8^ Studies have suggested that robustness can drive evolvability by buffering destabilising substitutions while permitting genetic variation to accumulate.^9^ This buffering is facilitated by chaperone-mediated assistance in folding, compensatory or suppressor mutations^10^, molecular overcrowding^11^, and oligomerization^12^. Since stability and function are largely dependent on three-dimensional structure, this prompts us to ask what makes some mutations deleterious while others neutral on the structural level, and which aspects of structure the neutrality can be attributed to.

The evolution of protein structures has been quantified and assessed through multiple mechanisms and has often been correlated with their structures.^13,14^ Early analyses demonstrated that buried residues evolve more slowly than surface-exposed residues, implicating solvent accessibility and packing as key determinants of evolutionary constraint.^15^ Subsequent studies showed that global structural properties such as protein length, contact density, secondary structure composition, and overall compactness correlate with evolutionary rate, often in highly interdependent ways^16,17^, thereby motivating the view that spatial architecture constrains sequence variation. It can thus be inferred that protein geometry, in addition to residue-level properties, shapes protein folds’ tolerance to mutation.

In this context, the concept of designability has been used in several studies as a unifying framework for these observed correlations. Theoretical work by Li^18,19^, Shakhnovich^19^, and colleagues proposed that highly designable structures are intrinsically more tolerant to mutation, as a larger fraction of sequence space maps onto the same fold. Bloom et al. describe how protein structure influences its rate of evolution, concluding that the rate of sequence evolution is attributed to the designability of the sequence, defined as the number of sequences that encode a particular fold. The designability, in turn, is influenced by several structural descriptors, such as subunit density (characterised by inter-residue contacts), protein length, secondary structure composition, and solvent accessibility, which are correlated with one another and with evolutionary rate. Experimental studies further demonstrated that proteins with greater thermodynamic stability exhibit increased mutational tolerance, supporting the notion that structural buffering underlies robustness.^4,20^

Another study by Riera et al. reports that age also influences the rate of evolution.^21^ They note that younger proteins, despite being less designable, tend to evolve faster than older proteins, indicating that robustness, function, and evolutionary context interact in complex ways. Investigating these relationships at scale has been challenging because straightforward quantification of robustness is difficult. These findings underscore the need for approaches that directly assess fold-level robustness, which reflects a convolution of structural, functional, and population-level constraints. The internal constraints of the structure are augmented by protein-protein interfaces, which may reduce conformational freedom, stabilise subunit organisation, and impose additional geometric constraints. Individual studies have highlighted the role of interfaces in modulating evolutionary pathways and in predicting the direction of evolution.^22,23^

In the present study, we quantify fold-level mutational robustness using family size, defined as the number of high-confidence structural homologs obtained from Foldseek^24^ searches against the AlphaFold database clustered at 50%. This gives us a fold-oriented measure of the sequence diversity compatible with a given three-dimensional architecture. We observe variability in family size, along with a comprehensive set of structural descriptors spanning size, intra-subunit packing, surface geometry, and non-covalent interactions, across protein homomeric complexes with varying oligomeric states. We present a systematic analysis of how protein architecture shapes mutational robustness and demonstrate that protein geometry defines a unified architectural framework that underlies mutational robustness and drives the evolutionary expansion of protein folds.

## Results

### Dataset and Family size

We curated datasets consisting of four classes of proteins – monomers, homodimers, homotrimers, and homotetramers from 3DComplex^25^ using annotations from QSbio^26^ containing 11077, 9184, 781, and 2149 protein structures, respectively. We select homomers owing to their high prevalence across organisms ^25,27^, and use monomers as a control to study the effect of oligomerisation on robustness. Homomerization, irrespective of the oligomeric state, is an abundant and advantageous phenomenon in nature due to the presence of repetitive elements, which can form supportive entities, as observed in cytoskeletal structures. It also creates multivalence, essential for protein binding ^28^.

We quantify mutational robustness at the structural level using the number of significant structural alignment hits returned by Foldseek searches against the AFDB50 database. We also compute a sequence-centric family size obtained by aligning sequences against the Uniref50 database using BLAST. Hits were filtered to contain high-quality matches using coverage, TM-score, and e-value cut-offs (see methods). Finally, we selected proteins present in both prokaryotes and eukaryotes, assuming that all proteins in the dataset have had sufficient time to evolve from prokaryotes to eukaryotes. This is done to remove any bias that may arise from differences in emergence time.

We observe that, in general, oligomers belong to larger families than monomers (Figure S1), consistent with the finding that oligomerisation confers several advantages on the protein subunit.^29,30^ These drive the reuse of homooligomeric folds in evolution. In contrast, while BLAST-derived sequence family size ranges from monomers to oligomers, it shows weaker separation among oligomeric states (Figure S1). This might suggest that while oligomerisation accompanies both sequence and structural diversification, the evolutionary signature is more accurately captured at the fold level. This remains consistent with protein structures being more conserved than their amino acid sequences, also evidenced by sequence family size being magnitudes higher than fold family size.

In terms of individual mutation-induced perturbations, they might be stabilised by compensatory contacts arising from constraints and stabilising interactions at subunit interfaces imposed by oligomeric assembly, thereby buffering the structural consequences of mutations. Such stabilisation could permit greater sequence divergence while maintaining the same overall fold, ultimately contributing to the larger structural families associated with oligomeric proteins.

### Protein size establishes an architectural basis for mutational robustness

The size of a protein subunit plays a determining role in the overall packing and stability. It enables the formation of subdomains, secondary structural and functional elements, increasing opportunities for more extensive tertiary interaction networks. Additionally, it has been shown to influence evolutionary rates in combination with subunit packing positively.^5^ We hypothesised that it would exhibit a similar relationship with fold-derived protein family size. With literature as precedent^31,32^, we classified structures into small (50-200 amino acids), medium (200-400 amino acids), and large (>400 amino acids) based on the length of a single subunit to account for dataset size and the substantial variability in chain length (coefficients of variation: monomers=0.62; homodimers=0.52; homotrimers=0.55; homotetramers=0.41). Consistent with the literature, protein size shows a moderate positive correlation with family size(Table S1), suggesting that larger proteins enable fold families to expand. Notably, the presence of interfaces appears advantageous, as it promotes the formation of favourable noncovalent bonds during oligomerisation. The additional geometric constraints enhance robustness, as evidenced by larger family sizes for oligomers than for monomers (Figure 1) with comparable subunit lengths. The resulting increase in the structures’ thermodynamic stability^33–35^ makes this a common evolutionary strategy.

**Figure 1.**
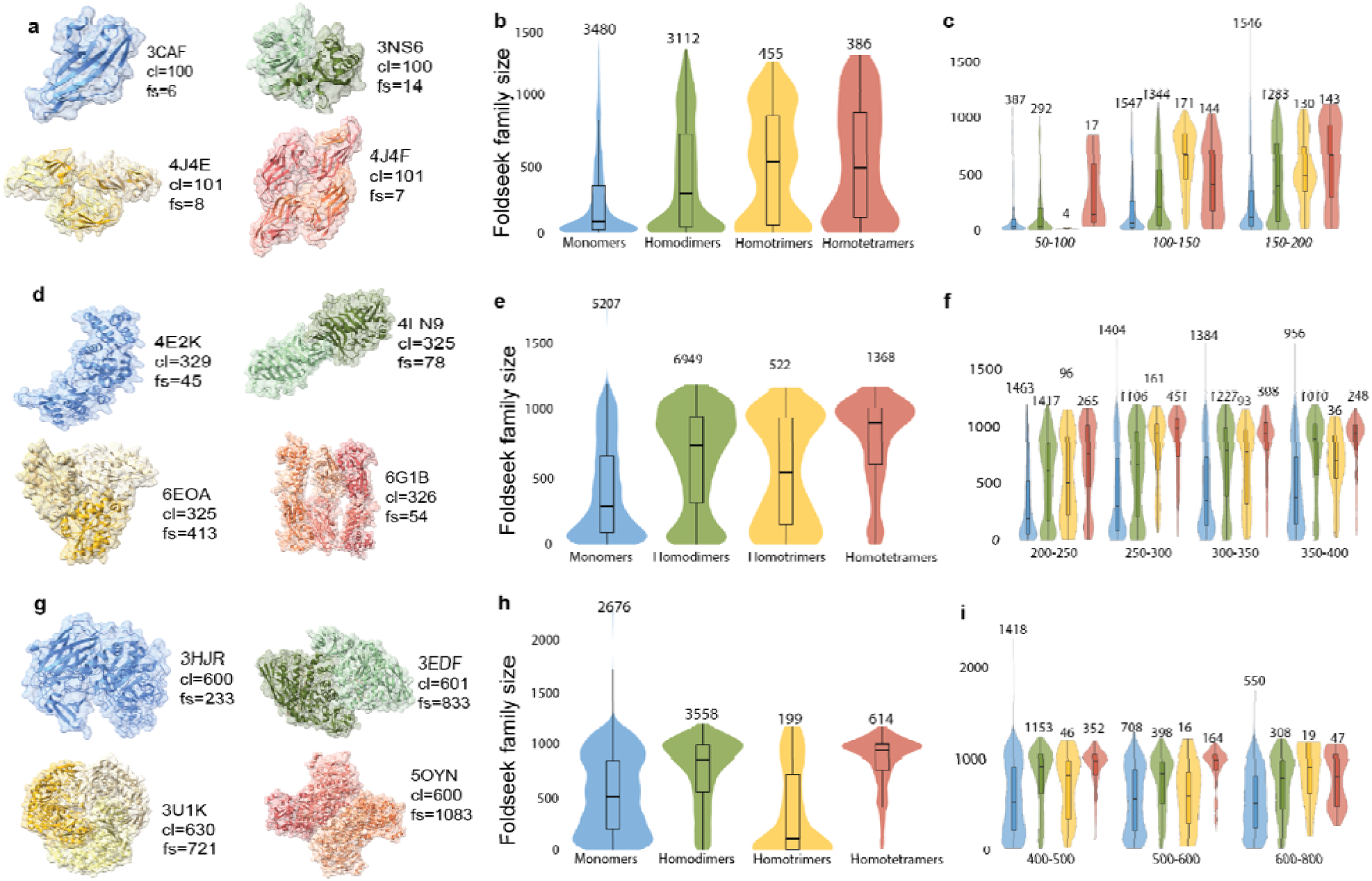
Variability in family size with increasing protein size. (a),(d),&(g) Representative structure of comparable sizes showing variation in family size with increasing degrees of oligomerisation. (b), (e) & (h). Higher-order oligomers exhibit a larger fold family size within size-stratified classes, except for trimers. A steady increase is observed particularly for subunits of size ∼500 amino acids(c),(f), and &(i). In very large subunits (>500 amino acids), family size decreases with increasing subunit size.

We further investigated the size-robustness relationship across proteins with varying functions, and found that the correlation holds across categories. As expected, functional categories differed markedly in their family-size distributions (Figure S2) and sizes. Enzymes, which constitute the largest functional class in our datasets (Figure S2), exhibited among the largest family sizes, particularly in higher oligomeric states (medians ranging from 750 to 1000), compared with monomeric enzymes (median = 341)(Table 1). They also tend to have larger subunits than proteins of the same oligomeric state that perform other functions. Consistent with their generally compact architectures and densely packed catalytic scaffolds. Previous studies have shown that enzyme folds are frequently reused throughout evolution and are often characterised by substantial mutational tolerance outside active-site regions. In contrast, signalling proteins exhibited the smallest family sizes across all oligomeric states (50-600) despite comparable chain lengths in some cases, suggesting that functional specificity and the requirement for finely tuned molecular recognition constrain sequence diversification. Regulatory and transport proteins occupied intermediate positions, whereas binding and structural proteins displayed broad distributions in family size and subunit size (Table S2), indicative of substantial heterogeneity in their underlying architectures and evolutionary histories.

**Table 1.**
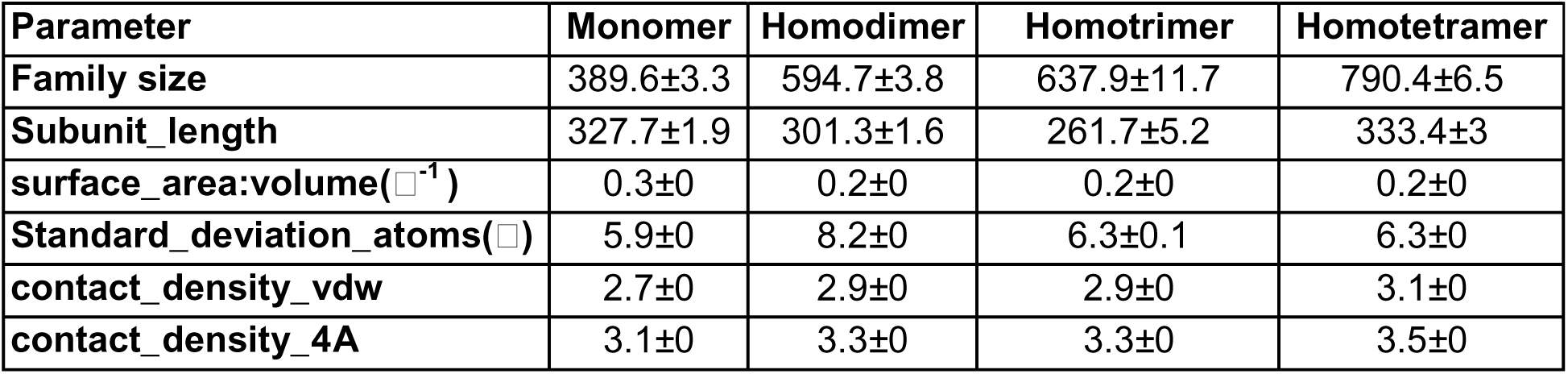
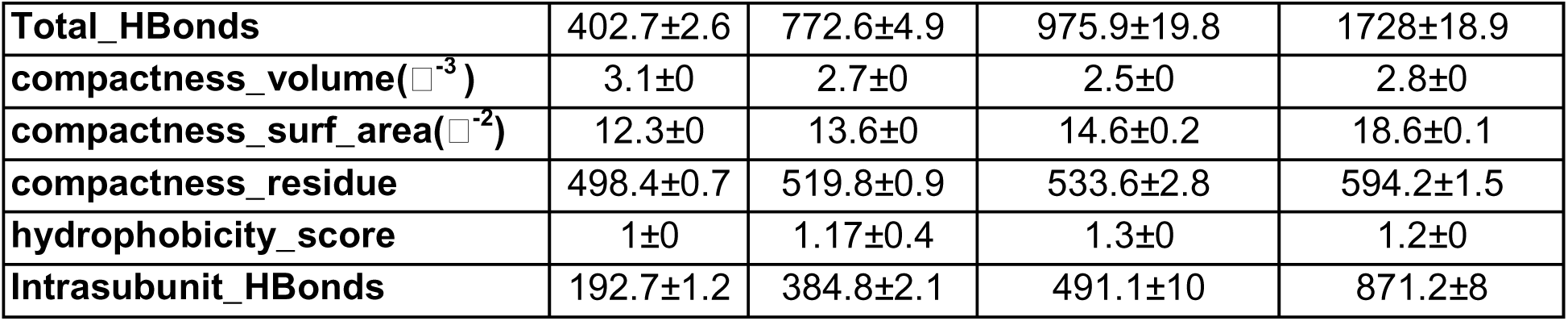
Summary of structural properties across homomeric proteins.

### Atomic packing scales with family size

The primary driver of protein folding is the need to fit within increasingly small spaces, such as cellular organelles and membranes. Additionally, there is the energetic imperative to minimise unfavourable solvent exposure. The efficient molecular packing of residues, driven and facilitated by size, amino acid composition, and dense contact networks formed by non-covalent interactions, is thus an important aspect of protein function as it affects the folding rate and stability.^36,37^ This led to the hypothesis that robustness would vary among proteins with different internal fold dynamics, and that quantifying it would reflect this relationship. We defined subunit packing spatially by measuring the number of residues in the protein core, normalising by volume and solvent-exposed surface area, assessing residue-residue and intra-subunit contacts, and measuring contact densities over a range of distances. Across oligomeric states, family size increases monotonically with measures of protein compactness (Figure 2). This trend is consistent across alternative methods of quantification(Figure S3). Subunit compactness correlates positively with family size in monomers (⍴∼0.5) and homodimers (⍴∼0.5) more strongly than in homotrimers (⍴∼0.3) and homotetramers (⍴∼0.3)(Table S1). It can be inferred that dense, intrasubunit contact networks are more robust to the destabilising effects of mutations. However, dependence on compactness and contact networks decreases with increasing subunit number, as this increases the opportunities and architectural space for buffering mutational consequences. The difference between a highly dense structure and a comparatively sparse protein is highlighted in Figures 2c and 2d. An N-citrylornithine decarboxylase (7KH2) has a large core (427 residues), contributing to a highly interconnected network and belonging to a larger structural family (891) than a metal-binding structural protein (3L1F) (family size = 37). This also supports our conclusion that enzymes, in general, belong to larger fold families and exhibit all structural features associated with greater robustness. This observation is further supported by the negative correlation between family size and solvent-exposed surface area per unit volume (Fig. 3), reiterating that proteins with greater solvent exposure and reduced geometric compactness are less tolerant of sequence variation.

**Figure 2.**
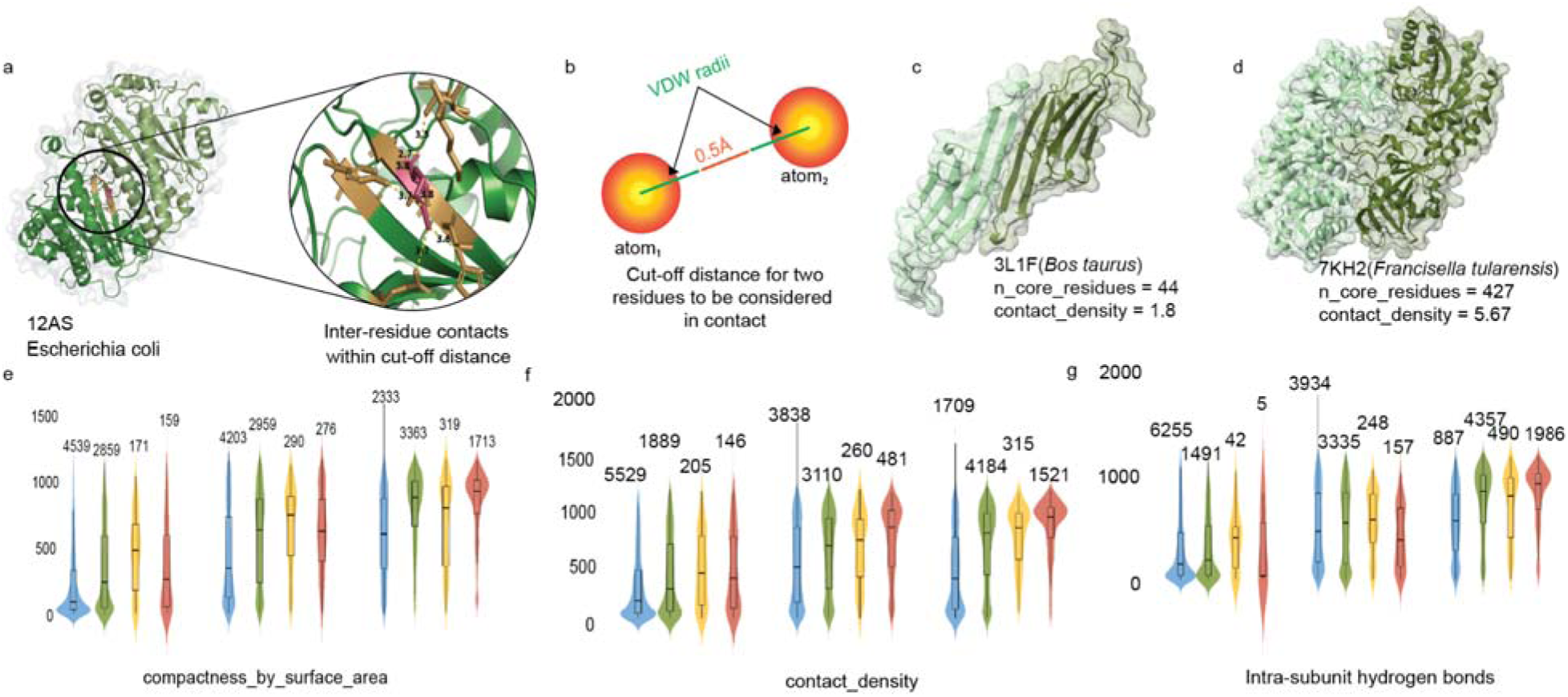
Family size scales with intrasubunit packing. (a) Schematic showing non-covalent interactions between amino acid residues within a subunit.(b) Cut-off distance for calculating the contact density of a subunit. (c),(d) Representative structures showing low and high contact density, characterised by the number of residues in the subunit core. (e) Family size increases with subunit compactness (defined as the number of residues in the core contained by a unit of solvent-exposed surface area) across oligomeric states. (f)Contact density (the average number of neighbouring residues that an amino acid interacts with) also positively influences family size. (g) Subunits with a denser hydrogen-bonding network have a larger family size.

**Figure 3.**
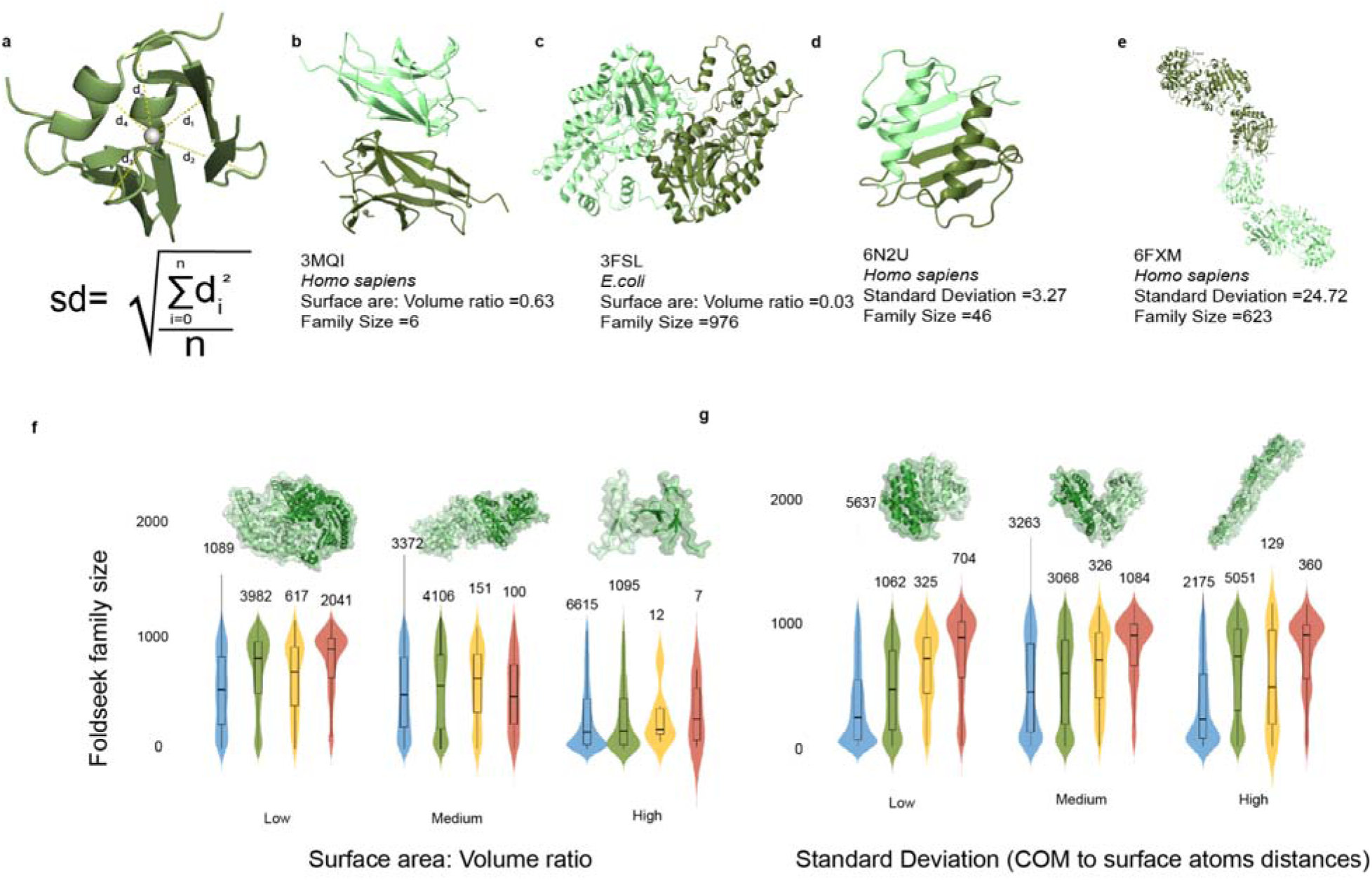
Correlation between surface geometry and robustness. (a) Quantification of surface geometry using the standard deviation of distances from the centre of mass to surface atoms.(b),(c) Representative structures showing high and low values of surface area to volume ratio and corresponding family sizes. (d), (e) Extreme values of standard deviation corresponding to globular(low standard deviation) and highly elongated (high standard deviation) structures exhibit markedly different family sizes. (f) Family size decreases with increasing surface-area-to-volume ratio. (g) Negative correlation of family size with standard deviation.

### Proteins’ robustness changes with shape and global surface geometry

The three-dimensional shape of the protein complex allows it to effectively perform its function, such as binding to ligands, nucleic acids, or contributing to larger multi-protein complexes that make up structural frameworks in the cell, implying that perturbations in protein shape due to mutations might lead to 3D structure collapse and possible disease or protein degradation by cellular machinery. This is evident in the widely studied mutation in haemoglobin from GLU to VAL, which distorts the globular (tetrameric) quaternary structure into an elongated shape driven by oligomerisation, causing sickle cell anemia^38^. This leads us to theorise that a protein’s shape imposes an essential constraint on structural alteration during evolutionary expansion. We thus examined the variability in protein family size with parameters representing shape. We quantified the shape of the protein using atomic distance measurements from its centre of mass to the solvent-accessible surface, according to the methods previously described by Mallik et al.^39^ An atom is considered to be on the surface if it has a solvent-accessible surface area of >= 5□^2^. Solvent accessibility of all atoms was calculated using NACCESS. The standard deviation of all atomic distances was used as a measure of globularity (Figure 3). The standard deviation for a globular/spherical protein would be closer to 0 than that of an elongated structure. We find a moderate positive correlation between shape heterogeneity and family size (Figure 3), suggesting that features such as local asymmetry or domain-like organisation may increase tolerance to sequence variation. In contrast to monomers and homodimers, homotrimers display a substantial negative correlation between shape irregularity and family size (⍴=-0.280), while homotetramers show an insignificant correlation (Table S1).

The variable relationship between global shape and mutational tolerance in different oligomeric states suggests that its effect is driven by architectural context. In monomers and dimers, the dependence on precise geometric symmetry is less than that of trimers and tetramers because there are fewer interface-mediated constraints, allowing local disruptions to be absorbed without destabilising QS. This suggests that moderate shape heterogeneity, reflecting asymmetry or domain protrusions, may enable structural flexibility or modular organisation, thereby increasing tolerance to sequence variation. In the case of higher-order oligomers, the constraints imposed by symmetry and cooperative interface packing limit the potential for surface irregularity, thereby preserving QS stability.

### Constraints from quaternary organisation enhance robustness

Our observations suggest a model of mutational robustness that is rooted in protein geometry, wherein architectural context is the primary driver of tolerance to sequence variation. While the spatial residue organisation within the subunit is key to compactness and subsequent contribution to family size expansion, the quaternary organisation of the subunits augments this property. The enhancement of mutational robustness by homooligomerization has been a ubiquitous observation and a central insight of this study. Oligomeric proteins consistently belong to larger structural families (Figure S1) and exhibit stronger geometry–robustness relationships than monomers of comparable size, emphasising the role of protein–protein interfaces in providing stabilising constraints that expand mutational tolerance^12^.

It is quite interesting to note that the correlation between structural properties such as subunit size, compactness and contact density and fold family size is lower in higher order homooligomers(trimers and tetramers) as compared to monomers and dimers(Table S1), and the median sizes and compactness remain comparable across monomers and oligomers(Table 1), the median family sizes increase remarkably with increasing oligomeric state(Table 1). This might suggest that, despite the reduction in overall conformational freedom induced by interfaces, oligomerisation constitutes an architectural extension of the monomeric unit’s qualities and reinforces the same principles that promote robustness in monomeric folds. This supports the findings of several studies, which find oligomerisation to be evolutionarily advantageous.^33,40,41^

Notably, throughout the study, we find that homotrimers do not follow the progression of observed parameters with increasing oligomeric state and does not exhibit the structure-robustness relationship as one might expect with respect to other categories examined here. This aligns with the general observations that homotrimers, or any oddmers for that matter, are underrepresented in the multimeric structure space^42^ and are relatively less preferred by evolution^22,42^ compared to proteins having even-numbered subunits. This might be attributed to homotrimers adopting a different strategy of oligomerization, wherein they have to utilise only heterologous interfaces to form a closed structure as opposed to isologous interfaces, which stabilize the multimeric form.^42^ The unique relationship of homotrimers along with higher order oddmers warrants further investigation.

### Functional necessities influence the extent of mutational robustness

To determine whether the relationship between structural organisation and robustness varies with protein function, proteins were grouped into broad categories based on Gene Ontology (GO) annotations before analysis. Enzymes constitute the largest class across datasets (53-71%), followed by other categories, including binding, regulatory, signalling, structural, and transport proteins (Figure S2). We hypothesised that proteins performing distinct functions would exhibit different relationships between architecture and family size, reflecting differences in evolutionary constraints.

Despite substantial differences in the absolute distribution of family size, the influence of oligomerisation was consistent across functional classes (Figure S2). Oligomeric proteins generally belonged to larger families than their monomeric counterparts and exhibited progressively higher compactness, contact densities, and lower solvent exposure per unit volume(Figure S3). Despite a conserved quaternary organisation-robustness relationship, the extent of the structure-robustness relationship varied markedly across functional categories. Enzymes consistently occupied the largest families and displayed the highest robustness measures across oligomeric states (Table S2). Family size and compactness increase threefold from monomeric to tetrameric enzymes, accompanied by a progressive reduction in surface exposure relative to volume. These observations are consistent with the extensive structural diversification of enzyme superfamilies, in which highly stable catalytic folds accommodate substantial sequence variation^9,43^. In contrast, proteins involved in signalling and regulatory functions consistently occupied among the smallest families, exhibiting lower compactness, reduced contact densities and greater solvent exposure than enzymes or binding proteins. These observations likely reflect stronger evolutionary pressures arising from the need to preserve high specificity of molecular recognition, transient protein-protein interactions, and allosteric communication, which are sensitive to sequence perturbation ^23^.

Transport proteins exhibit an interesting departure from the overall trend. Despite possessing the longest monomeric subunits and relatively large monomeric families, the increase in family size is not proportional to the degree of oligomerisation. They exhibit intermediate compactness and contact densities, indicating that geometric organisation, rather than protein size alone, is the stronger determinant of structural robustness.

These results demonstrate that biological function influences the magnitude of protein-fold diversification but not the structural descriptors that underlie their robustness. Proteins performing diverse functions find a common ground on architectural factors such as compactness, dense residue-contact networks, and reduced solvent exposure to achieve enhanced robustness, reiterating that structural geometry exerts a more fundamental influence on mutational robustness than functional necessities.

### Principal Component Analysis reveals the combined effect of structural determinants on robustness

We further wanted to assess whether the influence of structural factors on evolutionary fold expansion is collective rather than individual, since several properties point to the same global architectural quality in distinct ways. We correlated the properties and found significant correlations among groups of structural properties (Figure S4), indicating that they are interdependent to some extent and reflect a common underlying architectural framework. To this end, Principal Component Analysis was performed across oligomeric states using size, compactness, contact density, number of hydrogen bonds, surface geometry, and surface area: volume ratio.

Across oligomeric states, the first principal component (PC1) explained a significant portion of the total structural variation, accounting for ∼50% of the variance across datasets (Figure 4). Together, the first two principal components explained over 65% of the total variance (Figure S5). This indicates that the measured structural properties could be represented by a low-dimensional structural space. Inspection of PC1 loadings revealed that protein length, hydrogen bonding, compactness, and surface geometry contributed most strongly to the dominant principal axis. In contrast, contact density and surface area-to-volume ratio contributed comparatively less (Figure S5).

**Figure 4.**
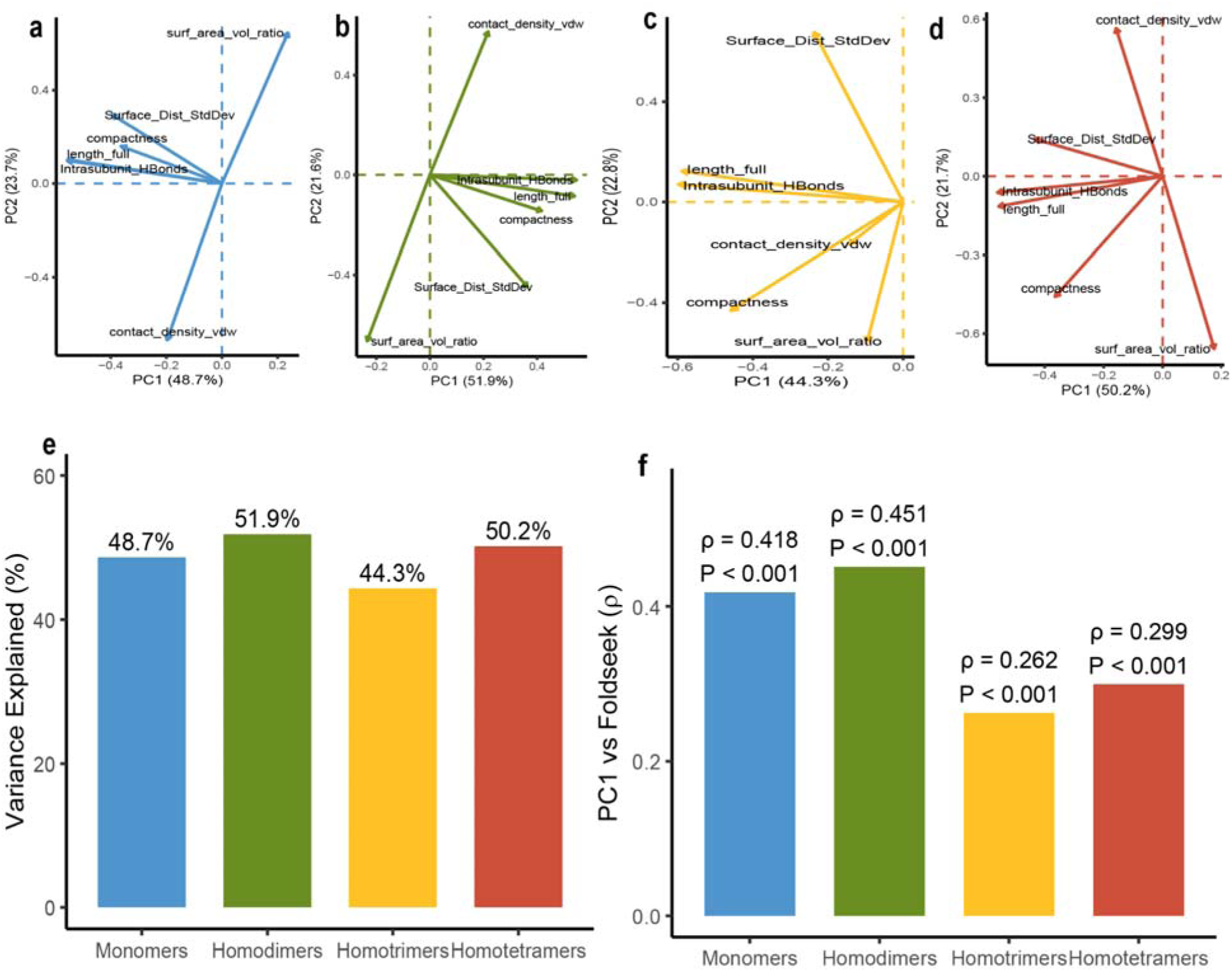
Principal component analysis of the structural determinants of robustness. (a)-(d) Contribution of structural descriptors to the first two principal axes. (e) Variance explained by the first principal axis(PC1) across oligomeric states. (f) Correlation between PC1 scores and foldseek-family size for each oligomeric state.

To investigate the association between the structural architecture and structural robustness, PC1 scores were correlated with foldseek family size (Figure 6). Significant positive correlations were observed for all oligomeric states, with the strongest correlation observed for homodimers (?=0.45) and monomers (? =0.42), followed by homotetramers (? =0.3) and homotrimers (? =0.26). As PC1 was primarily constituted by protein size, compactness, intrasubunit hydrogen bonds, and contact density, and an inverse contribution from solvent exposed surface area(Figure 4), the positive correlations suggest that proteins that form larger fold-families share a common architectural signature characterised by bigger subunits, dense residue packing, lower solvent exposure per unit volume, and greater overall constraints. It can be inferred that these structural qualities cooperate to define a geometric framework that governs the extent of mutational robustness.

## Discussion

Multiple structural descriptors that influence protein folding can tolerate sequence variation and diversify over time. The positive correlation between protein subunit size and family size suggests that larger folds are generally more compatible with a broader sequence space. This can be attributed to the availability of more secondary-structure elements and tertiary interaction motifs, which might increase the capacity for compensatory interactions or spatial adjustments that buffer destabilising mutations. Notably, subunits that are part of oligomeric complexes(dimers, trimers, and tetramers) exhibit enhanced robustness compared to monomers of comparable size. This observation suggests that size-induced tolerance is amplified by additional geometric constraints arising from subunit interfaces introduced by quaternary organisation. Importantly, this effect is non-trivial in the context of oligomeric proteins, where only single-subunit length was considered. Our results suggest that larger proteins exploit greater structural complexity to establish more densely connected residue networks that buffer the destabilising effects of mutations. Proteins of similar length can differ markedly in their tolerance to sequence variation, underscoring the importance of residue arrangement in three-dimensional space. Efficient packing, dense contact networks, and reduced surface exposure determine whether the robustness potential afforded by size is realised. The observation that oligomeric proteins exhibit greater robustness than monomers of comparable subunit length indicates that size-derived buffering is amplified by additional geometric constraints imposed by quaternary organisation, rather than by chain length alone.

A notable exception to these trends was observed among homotrimeric proteins, which frequently deviate from the relationships observed for monomers, dimers, and trimers, particularly in shape descriptors. Homotrimers possess a unique threefold symmetry, in which each subunit contributes to two equivalent interfaces, imposing geometric constraints distinct from those of dimers or tetramers. The influence of oligomerisation on mutational robustness may be further shaped by symmetry and the assembly pathway.

The structure-robustness relationship is consistent across functional classes despite substantial differences in absolute family sizes. Enzymes occupy the largest fold families and exhibit the highest compactness and packing densities. In contrast, signalling and regulatory proteins occupied comparatively smaller structure families, likely reflecting stronger constraints imposed by highly specific molecular recognition. This is essential owing to their shorter subunit lengths, which increase the pressure to conserve interacting residues and leave a smaller window for sequence diversification.

The PCA analysis extends the univariate correlation analyses by demonstrating that the global structural descriptors do not behave as independent variables but instead define a common axis of structural organisation. Protein length, compactness, intramolecular hydrogen bonding, and surface geometry consistently emerged as the dominant components across all oligomeric states, suggesting that proteins within larger Foldseek structural families share an integrated architectural framework rather than a single defining feature. The positive association between PC1 and Foldseek family size further indicates that structural family expansion is linked to the combined effects of these global structural characteristics. This observation strengthens the conclusion that mutational robustness is likely influenced by coordinated structural organisation, in which multiple interdependent features collectively contribute to a protein fold’s capacity to tolerate sequence divergence.

By directly quantifying robustness rooted in a structural context using fold-based family size, our study bridges concepts from protein evolution, structural biology, and designability theory. Our findings suggest that oligomerization should not be viewed simply as an increase in molecular complexity, but as an evolutionary strategy for expanding the accessible neutral sequence space. Although homo-oligomeric assemblies are thought to arise from monomeric ancestors through the acquisition of stabilising interfaces, the resulting increase in packing density and thermodynamic stability may allow these structures to tolerate a broader range of mutations, promoting the subsequent expansion of structural families. Conversely, the predominance of monomeric proteins in extant proteomes likely reflects the lower evolutionary and energetic costs of maintaining single-chain architectures rather than superior mutational robustness. Oligomerization therefore, represents a trade-off between architectural simplicity and evolutionary capacity, with interface formation enabling greater exploration of sequence space when increased robustness or functional complexity provides a selective advantage.

## Conclusion

Our findings suggest that, in general, mutational robustness in proteins is strongly influenced by the geometric organisation of the intrinsic fold. Proteins forming larger fold families are characterised by greater size, higher intra-subunit packing, denser residue-residue contact networks, and stabilising inter-subunit interactions in oligomers. These qualities collectively increase the structure’s capacity to accommodate sequence variation while maintaining structural integrity. Additionally, we observed differences across functional classes, suggesting that evolutionary pressures shape robustness in ways that reflect functional requirements. Further studies on the impact of oligomerisation-imposed constraints will provide new insights.

## Methods

### Dataset assembly and oligomer classification

Protein structure datasets were compiled separately for monomers, homodimers, homotrimers, and homotetramers from experimentally determined PDB structures from 3DComplex^25^. The datasets are non-redundant up to a level of 90% and have a QSbio annotation of “NO” (representing physiologically relevant, high-confidence structures with no QS annotation error from PDB) or “PROBNOT”(representing physiologically relevant, medium-confidence structure with probably no QS annotation error in PDB), and/or QSbio error probability <5%; we relied on the error probability solely when no annotation is present for a structure entry. Only biologically relevant assemblies annotated as homomeric were included. For oligomeric proteins, analyses were performed at the level of a single representative subunit to avoid trivial scaling effects due to the total complex size. Structures with unresolved regions, high model error, or ambiguous quaternary assignments were excluded based on quality filters (pdb_error, pdb_error2, QSbio_err_probability).

Each dataset was curated to remove redundancy and retain one representative per structural entity. Symmetry assignments were recorded from biological assembly annotations.

### Family size estimation using Foldseek and BLAST

Fold-level mutational robustness was quantified using family size, defined as the number of significant structural matches returned by Foldseek searches against the AFDB50 database. Foldseek aligns protein structures by encoding tertiary residue–residue interaction patterns as sequences over a structural alphabet and then performing rapid structural alignment in this reduced-representation space.

Each query structure was searched against AFDB50 using default alignment parameters. Significant matches were filtered using alignment coverage and E-value thresholds that were consistent across all queries. We filtered the hits, selecting TMscore >= 0.65, coverage of >=80%, and e-value <0.05 to obtain a Foldseek family size, reflecting a fold-centric robustness metric that assesses the size of sequence space compatible with a given fold. The total number of retained matches was recorded as the Foldseek family size for each structure. Because Foldseek alignment is independent of amino acid sequence identity, this metric reflects the diversity of sequences compatible with a given three-dimensional architecture rather than direct sequence similarity. The generated homologs were filtered for per cent query coverage (>=80%) and e-value(<=0.01). For both family-size metrics, we further filtered the homologs for copy number (taking one sequence per organism) to remove bias introduced by gene duplication events.

### Contact density and compactness

Residue-residue contacts were defined as heavy-atom contacts within specified distance thresholds. Contacts were counted when any pair of heavy atoms from residues *i* and *j* were within (a) 3.5, (b) 4.0, (c) 4.5, and (d) the sum of VDW radii of the atom pair +0.5 of each other. Consecutive neighbour contacts were excluded. The number of contacts was normalised by the number of residues in the subunit to get “contact density”.

Compactness was defined by the bulk of the protein core contained within the unit volume, unit solvent-exposed surface area, and the fraction of total residues residing in the core. Structural location of residues based on solvent accessibility was defined as described previously by Levy^44^. Accessible surface areas were calculated using NACCESS. Core residues were defined as those having relative solvent accessibility < 25%.

### Quantification of global surface geometry

The overall shape was defined based on surface irregularities. Distances from the structure’s centre of mass to surface atoms were measured, and their standard deviation was calculated. This metric was used to quantify the shape based on the rationale that the standard deviation would be closer to 0 for a more globular protein.

### Polar and non-polar interactions

Hydrogen bond information was generated using HBPLUS^45^ and classified as intra-subunit, inter-subunit, or water-mediated based on the origin of the donor and acceptor atoms. Non-heavy atoms and hydrogen bonds between adjacent residues were ignored during the calculation.

### Functional Annotation of Proteins

Assigned functions of proteins were obtained using the Gene Ontology(GO) tool^46^, followed by broader functional assignment via keyword search with an in-house Python script. GO annotations containing the words “catalytic activity” or the suffix “ase” were classified as enzymes; annotations containing “structural”, “transport”, or “binding” were classified as such.

### Statistical analysis

All statistical analyses were performed in Python using pandas, NumPy, and SciPy. Associations between structural parameters and Foldseek family size were assessed using Spearman’s rank correlation coefficient to capture monotonic relationships independent of linearity assumptions. Comparisons across oligomeric states were performed independently for each dataset. Distributional differences across parameter tertiles were visualised using violin and box plots using R^47^. Where appropriate, size-controlled comparisons were conducted by matching proteins within narrow length windows to evaluate geometry-dependent effects independent of chain length.

## CRediT authorship contribution statement

**Srijita Acharya:** Writing – original draft, Formal analysis, Data curation, Methodology, Investigation. **Sucharita Dey:** Writing – review & editing, Resources, Project administration, Funding acquisition, Conceptualisation, Supervision.

## Acknowledgments

SA acknowledges the MHRD fellowship. SD acknowledges DBT India for the RLS grant (RLS grant to SD: BT/RLF/Re-entry/10/2020, sanction order serial number 145), ARG MATRICS grant from ANRF (ANRF/ARGM/2025/000752/TS) and IIT Jodhpur, India, for infrastructure support.

## Data availability

Data will be made available on request.

