## Supplementary file for "Protein size and geometry govern mutational robustness"

**Table S1. Statistical analysis of the correlation between structural parameters and FoldSeek family size, including Spearman Correlation coefficients and p-values.**

| **Parameter** | **Spearman_rho** | **p_value** |
| --- | --- | --- |
| **Monomers** | | |
| length_full | 0.40 | 0 |
| compactness_by_surf_area | 0.50 | 0 |
| compactness_by_residue | 0.50 | 0 |
| compactness_by_volume | 0.42 | 0 |
| contact_density_3.5A | 0.28 | 7.9E-194 |
| contact_density_4A | 0.39 | 0 |
| contact_density_vdw | 0.31 | 8.96E-242 |
| surf_area_vol_ratio | -0.42 | 0 |
| Surface_Dist_StdDev | 0.12 | 3.55E-34 |
| hydrophobicity_by_index | -0.11 | 2.94E-30 |
| Total_HBonds | 0.38 | 0 |
| Intrasubunit_HBonds | 0.39 | 0 |
| **Homodimers** | | |
| length_full | 0.42 | 0 |
| compactness_by_surf_area | 0.54 | 0 |
| compactness_by_residue | 0.52 | 0 |
| compactness_by_volume | 0.44 | 0 |
| contact_density_3.5A | 0.17 | 3.01E-58 |
| contact_density_4A | 0.35 | 1.55E-259 |
| contact_density_vdw | 0.25 | 8.78E-132 |
| surf_area_vol_ratio | -0.37 | 1.67E-301 |
| Surface_Dist_StdDev | 0.14 | 9.37E-44 |
| hydrophobicity_by_index | -0.09 | 2.42E-18 |
| Total_HBonds | 0.45 | 0 |
| Intrasubunit_HBonds | 0.46 | 0 |
| **Homotrimers** | | |
| length_full | 0.19 | 5.20E-08 |
| compactness_by_surf_area | 0.26 | 1.40E-13 |
| compactness_by_residue | 0.29 | 4.72E-16 |
| compactness_by_volume | 0.28 | 7.59E-16 |
| contact_density_3.5A | 0.34 | 5.09E-23 |
| contact_density_4A | 0.34 | 2.04E-22 |
| contact_density_vdw | 0.32 | 2.07E-20 |
| surf_area_vol_ratio | 0.03 | 0.457718 |
| Surface_Dist_StdDev | -0.04 | 0.271882 |
| hydrophobicity_by_index | 0.12 | 0.000539 |
| Total_HBonds | 0.28 | 1.84E-15 |
| Intrasubunit_HBonds | 0.29 | 8.46E-17 |
| **Homotetramers** | | |
| length_full | 0.27 | 2.10E-37 |
| compactness_by_surf_area | 0.37 | 1.41E-71 |
| compactness_by_residue | 0.36 | 1.25E-67 |
| compactness_by_volume | 0.28 | 5.20E-39 |
| contact_density_3.5A | 0.34 | 1.58E-59 |
| contact_density_4A | 0.29 | 1.35E-41 |
| contact_density_vdw | 0.23 | 2.05E-27 |
| surf_area_vol_ratio | -0.21 | 1.07E-23 |
| Surface_Dist_StdDev | 0.04 | 0.088503 |
| hydrophobicity_by_index | 0.11 | 2.00E-07 |
| Total_HBonds | 0.36 | 3.18E-66 |
| Intrasubunit_HBonds | 0.35 | 1.98E-61 |

**Table S2. Median values of family size and structural parameters in different functional groups**

| **Structural feature** | **Monomers** | **Homodimers** | **Homotrimers** | **Homotetramers** |
| --- | --- | --- | --- | --- |
| **Enzymes** | | | | |
| Compactness | 0.49 | 0.87 | 1.11 | 1.65 |
| Surface_area:  volume ratio | 0.25 | 0.20 | 0.17 | 0.15 |
| Standard Deviation | 5.38 | 8.09 | 5.72 | 6.10 |
| foldseek_family_size | 341 | 781 | 758 | 927 |
| Subunit length | 308 | 316 | 258.5 | 332 |
| Contact density | 2.69 | 2.94 | 2.89 | 3.12 |
| **Regulatory Proteins** | | | | |
| Compactness | 0.41 | 0.62 | 0.52 | 1.01 |
| Surface_area:  volume ratio | 0.30 | 0.22 | 0.16 | 0.17 |
| Standard Deviation | 5.22 | 7.09 | 5.28 | 5.60 |
| foldseek_family_size | 80 | 296 | 479 | 273 |
| Subunit length | 208 | 194 | 115 | 177 |
| Contact density | 2.59 | 2.96 | 2.64 | 3.06 |
| **Binding Proteins** | | | | |
| Compactness | 0.45 | 0.73 | 0.86 | 1.58 |
| Surface_area:volume ratio | 0.27 | 0.21 | 0.15 | 0.16 |
| Standard Deviation | 4.95 | 7.48 | 5.33 | 5.37 |
| foldseek_family_size | 212 | 348 | 464 | 399 |
| Subunit length | 251 | 216 | 173 | 233 |
| Contact density | 2.65 | 2.78 | 2.77 | 2.85 |
| **Signalling Proteins** | | | | |
| Compactness | 0.43 | 0.66 | 0.83 | 1.30 |
| Surface_area:volume ratio | 0.29 | 0.24 | 0.14 | 0.18 |
| Standard Deviation | 5.65 | 7.44 | 5.52 | 5.36 |
| foldseek_family_size | 81.5 | 58.5 | 140 | 616.5 |
| Subunit length | 227.5 | 178.5 | 227 | 221 |
| Contact density | 2.52 | 2.64 | 2.61 | 3.15 |
| **Structural Proteins** | | | | |
| Compactness | 0.42 | 0.58 | 1.25 | 1.52 |
| Surface_area:volume ratio | 0.30 | 0.24 | 0.12 | 0.16 |
| Standard Deviation | 4.90 | 9.13 | 7.51 | 8.88 |
| foldseek_family_size | 100 | 109 | 231 | 686 |
| Subunit length | 200 | 207 | 436 | 387 |
| Contact density | 2.55 | 2.64 | 2.58 | 3.11 |
| **Transport Proteins** | | | | |
| Compactness | 0.48 | 0.71 | 1.23 | 1.29 |
| Surface_area:volume ratio | 0.25 | 0.22 | 0.17 | 0.16 |
| Standard Deviation | 6.15 | 8.27 | 6.91 | 6.29 |
| foldseek_family_size | 606 | 198 | 650 | 453 |
| Subunit length | 355 | 275 | 399 | 256 |
| Contact density | 2.75 | 2.92 | 3.24 | 3.21 |

**Table S3. Median values of family size and structural parameters across size-stratified datasets**

|  | **Monomers** | | | **Homodimers** | | | **Homotrimers** | | | **Homotetramers** | | |
| --- | --- | --- | --- | --- | --- | --- | --- | --- | --- | --- | --- | --- |
| **Parameter** | **Small** | **Medium** | **Large** | **Small** | **Medium** | **Large** | **Small** | **Medium** | **Large** | **Small** | **Medium** | **Large** |
| **Family size** | 95±4.59 | 308±4.89 | 528±7.09 | 302±6.37 | 742±5.23 | 864±7.26 | 580±16.34 | 801±17.39 | 780±39.22 | 515.5±19.42 | 921.5±8.02 | 956±8.91 |
| **Intrasubunit_HBonds** | 84±0.56 | 175±0.76 | 306±1.71 | 180±0.97 | 376±1.29 | 617.5±3.15 | 252±3.97 | 546±5.68 | 968±21.61 | 358.5±5.64 | 768±4.39 | 1278±8.43 |
| **Standard_deviation_atoms** | 3.7±0.03 | 5.27±0.03 | 7.24±0.05 | 5.97±0.03 | 8.02±0.03 | 9.63±0.07 | 5.11±0.07 | 5.88±0.12 | 7.36±0.5 | 4.91±0.07 | 5.7±0.03 | 7.27±0.07 |
| **Subunit_length** | 145±0.58 | 292±0.78 | 495±2 | 147±0.6 | 294±0.86 | 476±2.24 | 145±1.52 | 277±2.42 | 479±12.15 | 148.5±1.81 | 290±1.48 | 483±3.07 |
| **Total_HBonds** | 190±1.58 | 370±2.25 | 599±5.58 | 378±2.97 | 736±4.29 | 1228±11.88 | 601±11.94 | 1048.5±19.59 | 1616±82.15 | 747.5±16.56 | 1532±15.15 | 2454±39.77 |
| **compactness_surf_area** | 8.65±0.04 | 12.57±0.04 | 15.03±0.06 | 9.06±0.06 | 14.46±0.05 | 17.97±0.09 | 11.58±0.17 | 15.88±0.19 | 17.8±0.48 | 11.95±0.28 | 18.56±0.09 | 22.32±0.15 |
| **compactness_volume** | 2.6±0.01 | 3.18±0.01 | 3.49±0.01 | 1.99±0.01 | 2.92±0.01 | 3.22±0.01 | 1.85±0.03 | 2.87±0.03 | 3.21±0.06 | 2.06±0.03 | 2.82±0.01 | 3.21±0.02 |
| **contact_density_4A** | 2.89±0 | 3.12±0 | 3.2±0 | 3.14±0.01 | 3.37±0 | 3.43±0.01 | 3.21±0.03 | 3.41±0.02 | 3.4±0.03 | 3.31±0.02 | 3.53±0.01 | 3.59±0.02 |
| **contact_density_vdw** | 2.49±0.01 | 2.69±0 | 2.75±0.01 | 2.74±0.01 | 2.95±0 | 2.99±0.01 | 2.78±0.03 | 2.94±0.02 | 2.91±0.04 | 2.89±0.02 | 3.11±0.01 | 3.15±0.02 |

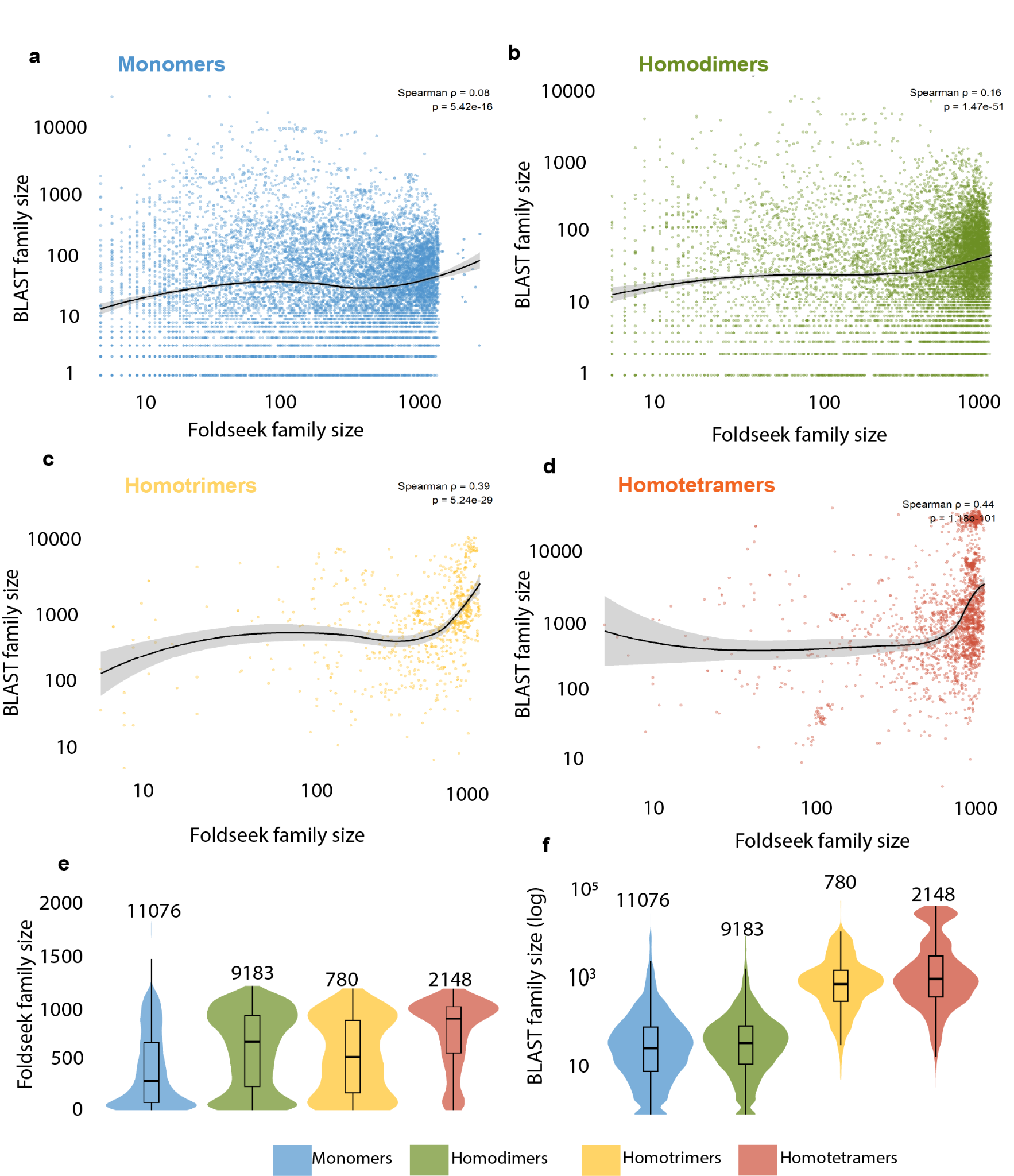

**Figure S1.Relationship between sequence- and structure-derived protein family sizes across oligomeric states.a–d**, Scatter plots comparing Foldseek structural family size with BLAST sequence family size for monomers (**a**), homodimers (**b**), homotrimers (**c**) and homotetramers (**d**). Black curves represent locally weighted regression (LOESS) fits. Spearman correlation coefficients (ρ) and corresponding *P* values are indicated.**e,**Distribution of Foldseek structural family sizes across oligomeric states. Numbers above violins indicate the number of proteins analyzed in each dataset.**f**, Distribution of BLAST sequence family sizes (log scale) across oligomeric states. Boxes indicate median and interquartile range.

**Figure S2. Structural determinants of protein robustness are conserved across functional classes. a** Distribution of Foldseek structural family size across functional classes for monomers (blue), homodimers (green), homotrimers (yellow) and homotetramers (red). Enzymes generally belong to the largest structural families, whereas regulatory and signalling proteins belong to comparatively smaller families.**b–f,** Distribution of structural descriptors across functional classes and oligomeric states, including subunit length **(b)**, surface area-to-volume ratio **(c)**, compactness **(d)**, residue contact density **(e)**, and surface irregularity (surface distance standard deviation) **(f)**. Across all functional classes, increasing oligomeric complexity is accompanied by greater compactness, reduced solvent exposure, and denser residue packing, indicating that the structural features associated with large structural families are broadly conserved irrespective of protein function. Box plots show the median, interquartile range, and 1.5× interquartile range; individual points represent outliers.
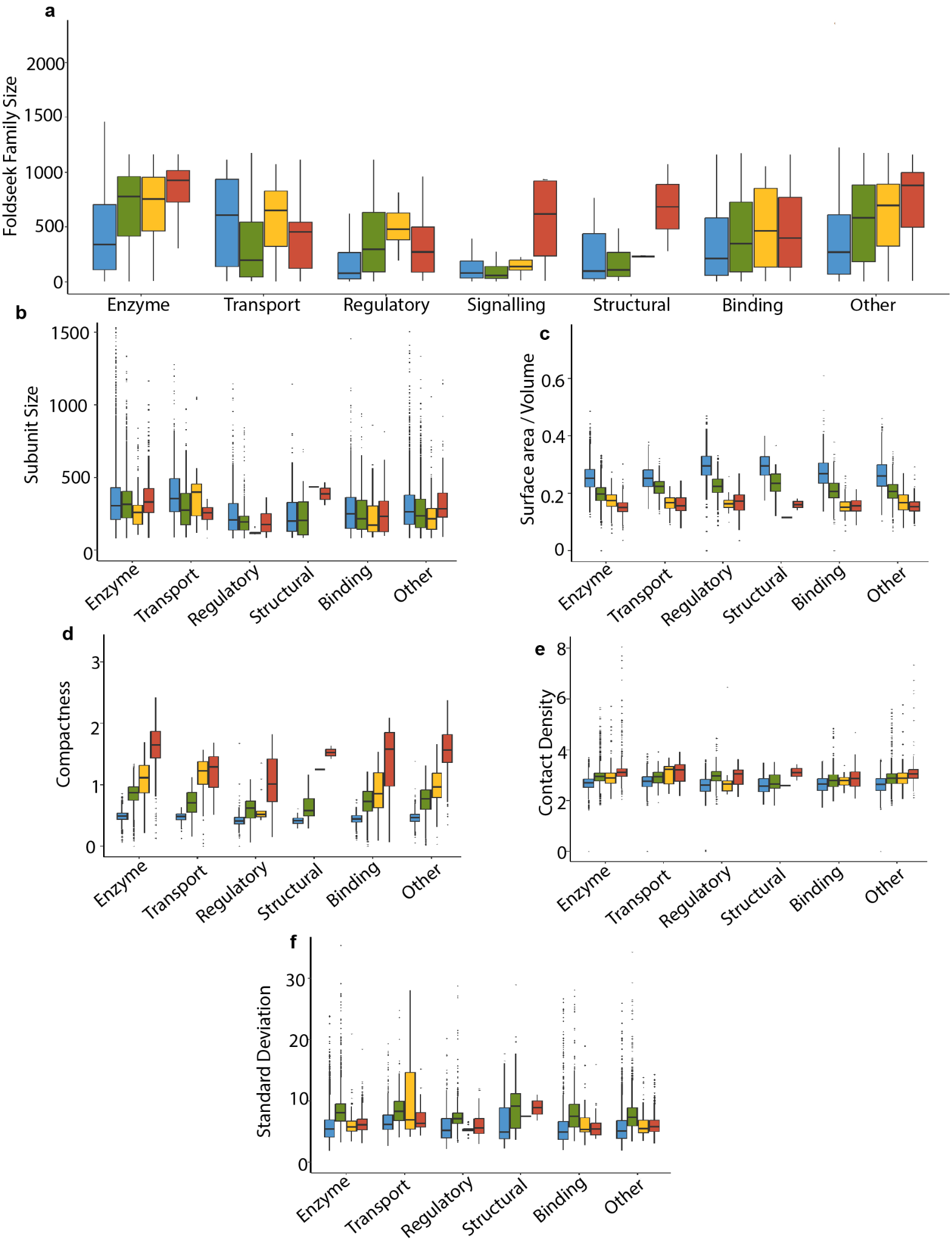

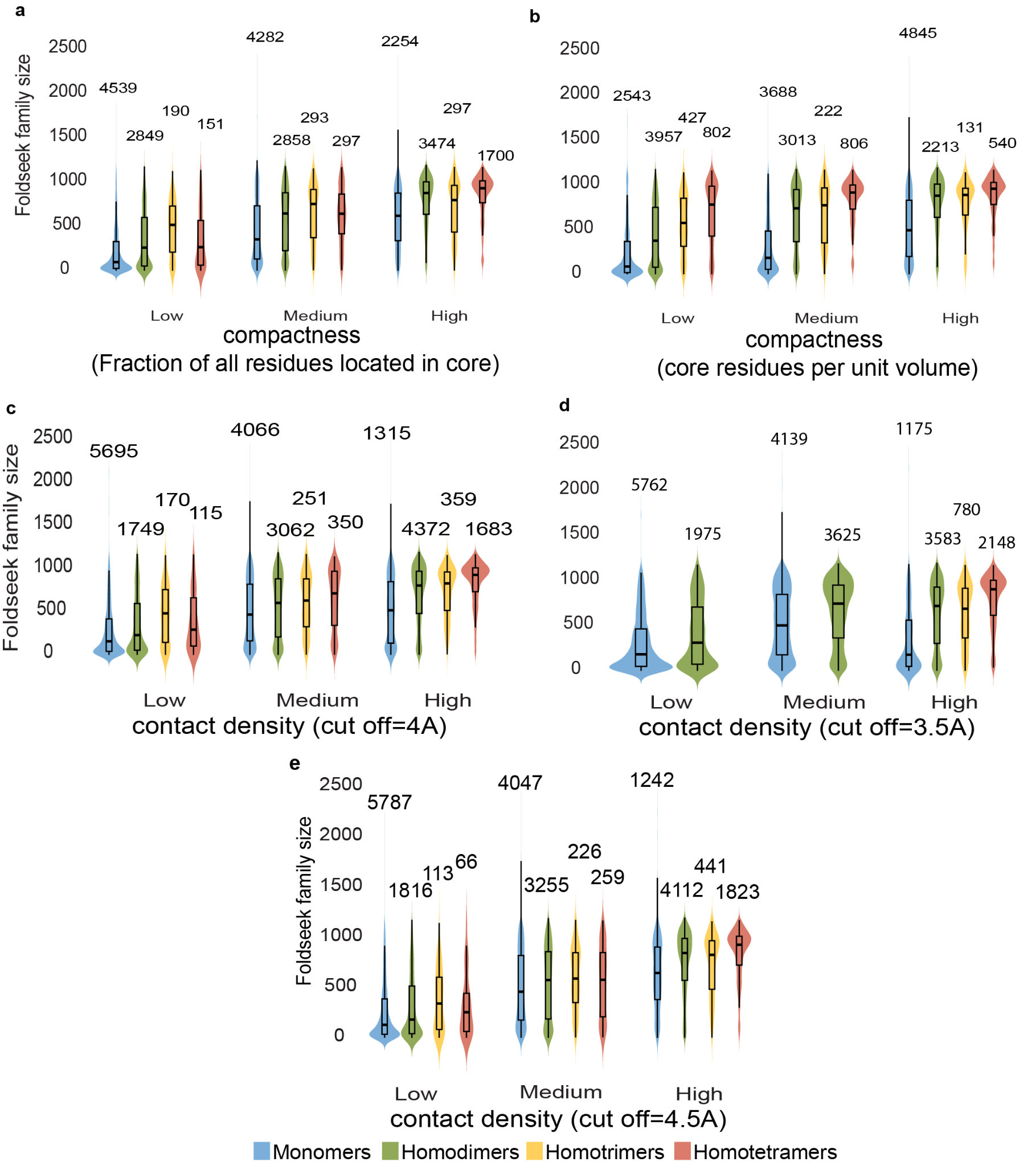

**Figure S3.Structural family size increases with multiple independent measures of protein compactness and residue packing.** Proteins were divided into low, medium and high groups using global tertiles for each structural descriptor. **a**, Fraction of residues located in the protein core. **b**, Core residue density normalised by protein volume.**c**, Residue contact density calculated using a 4Å distance cutoff. **d**, Residue contact density calculated using a 3.5 Å distance cutoff. **e**, Residue contact density calculated using a 4.5Å distance cutoff.For each panel, Foldseek family size distributions are shown for monomers, homodimers, homotrimers and homotetramers. Numbers above violins indicate the number of proteins in each bin.

**Figure S4.Pairwise correlations among structural descriptors associated with protein robustness.**Correlation matrices showing Spearman correlation coefficients between structural descriptors for **a**. Monomers, **b**. Homodimers, **c.** homotrimers**, d.** and homotetramers **(d)**. Positive and negative correlations are shown in red and blue, respectively. Only statistically significant correlations are displayed.
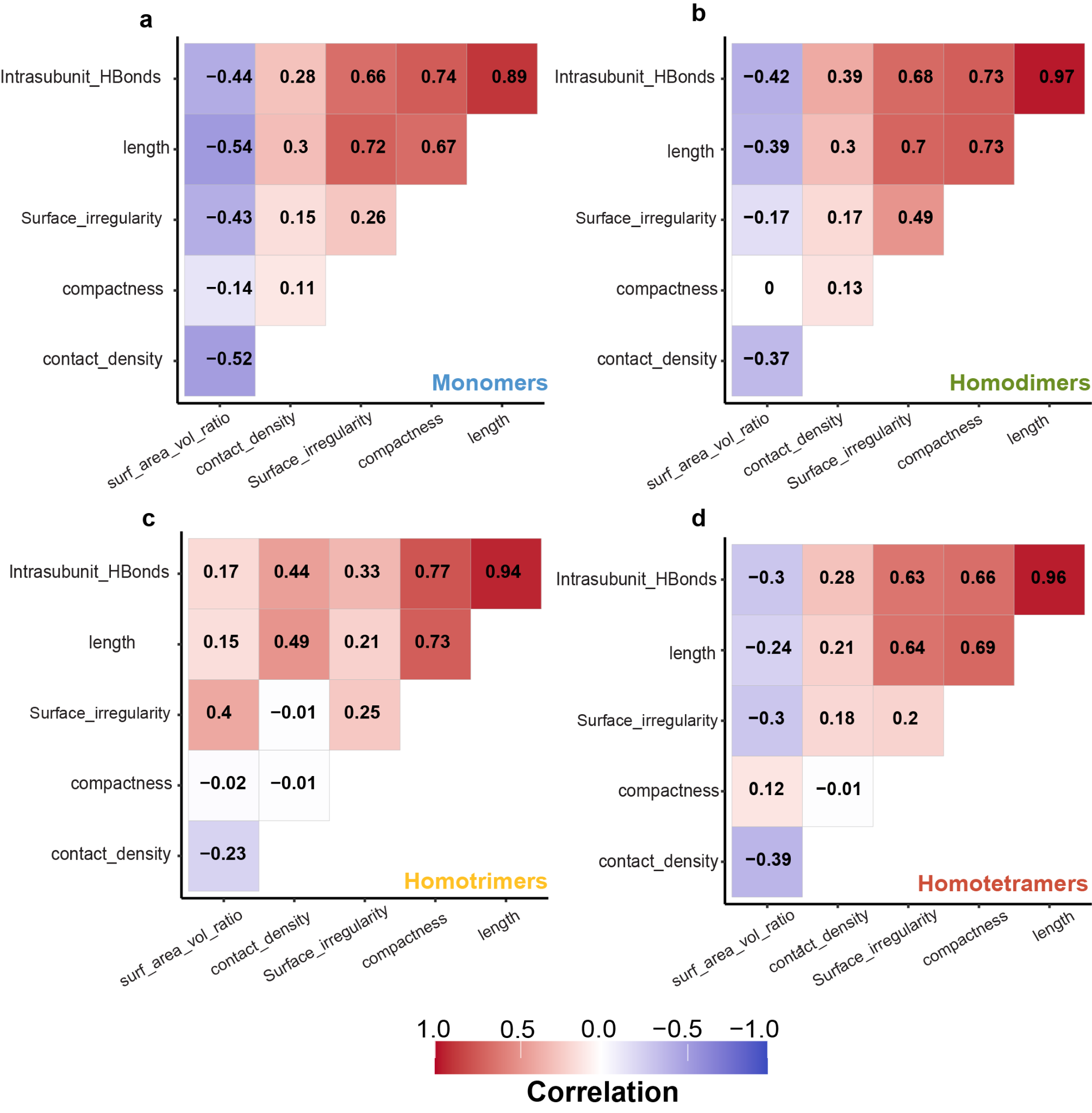

**Figure S5. Principal component analysis identifies the dominant sources of structural variation across oligomeric states.a–d**, Scree plots showing the percentage of variance explained by each principal component for monomers (**a**), homodimers (**b**), homotrimers (**c**) and homotetramers (**d**).**e**, Relative contribution of individual structural descriptors to the first principal component in each oligomeric state. Values represent the percentage contribution of each variable to PC1.
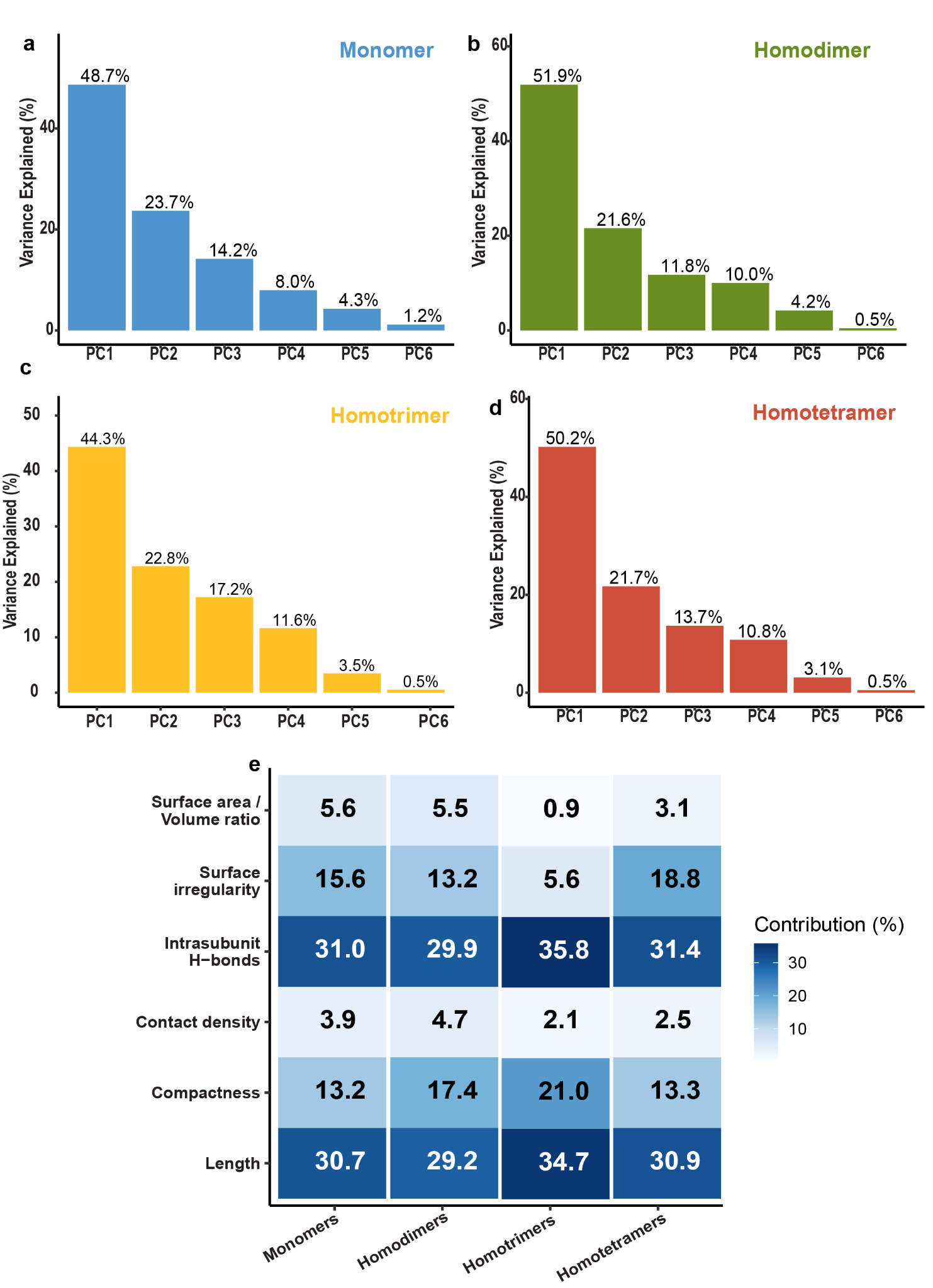

**Figure S6. Relationship between the principal structural axis and Foldseek family size.**Scatter plots showing Foldseek structural family size as a function of the first principal component (PC1) for **monomers (Spearman 𝜌 = 0.42) (a)**, **homodimers (Spearman 𝜌 = 0.45) (b)**, **homotrimers (Spearman 𝜌 = 0.26) (c)** and **homotetramers (Spearman 𝜌 = 0.3) (d)**. Solid lines indicate linear regression fits with 95% confidence intervals (grey shading). PC1 primarily reflects variation in protein size, compactness, residue packing and intramolecular hydrogen bonding, demonstrating that the dominant structural axis identified by principal component analysis is positively associated with structural family size across all oligomeric states.
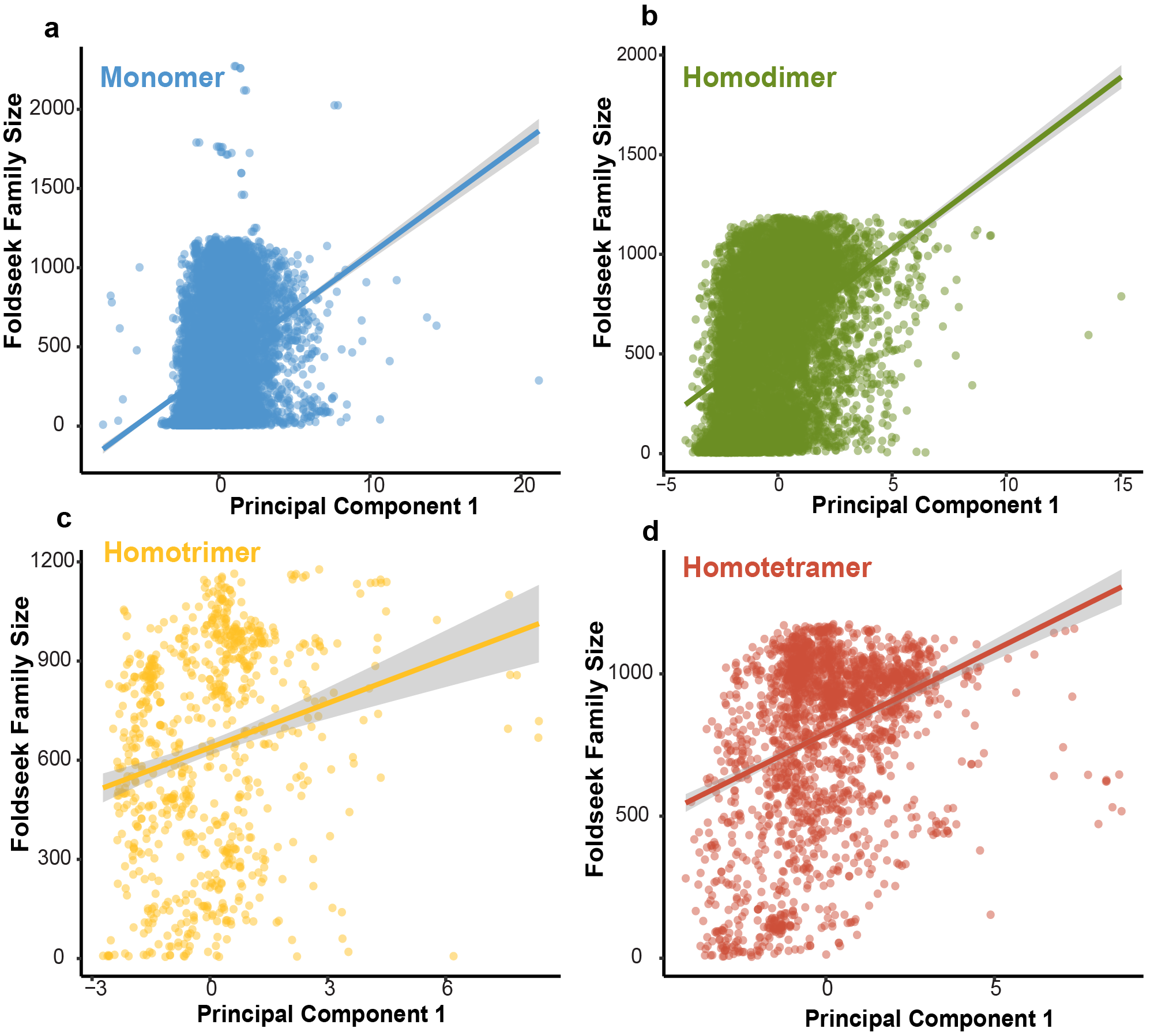
